# Funmap2: an R package for QTL mapping using longitudinal phenotypes

**DOI:** 10.1101/231928

**Authors:** Nating Wang, Tinyi Chu, Jiangtao Luo, Rongling Wu, Zhong Wang

## Abstract

QTL mapping is a powerful tool to infer the complexity of the genetic architecture underlying phenotypic traits, and has been extended to include longitudinal traits measured at multiple temporal/spatial points. Here, we introduce the R package *Funmap2* based on the functional mapping framework, which integrates biological prior knowledge into the statistical model. Specifically, the functional mapping framework is engineered to include longitudinal curves that describes the genetic effects, and the covariance matrix of the trait of interest. *Funmap2* may automatically choose the type of longitudinal curve and covariance matrix by information criterion. *Funmap2* is available for download at https://github.com/wzhy2000/Funmap2.

## Introduction

The advance in molecular biology has dramatically increased the number of molecular markers available to reveal the genome structure and organization of any organism. QTL mapping exploits these markers to identify the genomic regions associated with the quantitative traits within an inbred population. In the past 20 years, a variety of statistical models have been developed to detect QTLs, greatly facilitating the narrowing down of the genomic regions that control the biological traits. In addition to the study of additive and dominant effects, QTL mapping has been successfully applied to the epistasis effects, allometric growth and pleiotropic effects.

Lander and Botstein (Lander 1989) established tractable statistical methodologies to map QTL on one chromosomal interval bracketed by two flanking markers, known as interval mapping method. Later composite interval mapping improved interval mapping by including markers from other intervals as covariates to control the overall genetic background (Jiang, and Zeng, 1995). In 1999, Kao et al. proposed the simultaneous use of multiple marker intervals to map multiple QTL of epistatic interactions throughout a linkage map. Since then, applications of QTLs mapping in complicated genetic and genomic problems have boomed.

Albeit the advance in mapping resolution and extension to more complicated mapping problems, conventional mapping approaches are restricted to phenotypic data measured at a single time point. In many important biological problems, however, genotypes that control longitudinal traits, such as those measured during developmental process and environmental changes, cannot be effectively accommodated under the framework of single trait QTL mapping. Several approaches (Ma et al. 2002, Wu et al. 2002, Yang et al. 2006) have been developed for QTL mapping on such traits. Functional mapping (Ma et al. 2002, Wu and Lin 2006, Sun et al. 2015) is a statistical framework derived to map genes that control the dynamic biological process of complex traits. In this framework, a mixture model is fitted using EM algorithm through maximizing likelihood, followed by hypothesis testing the significance of association. In addition, model parameters describing growth trajectories can also be estimated. Functional Mapping has shown remarkable performance in associating QTLs with dynamic traits in plant (Zhao et al. 2004b, Li et al. 2010b, Yang et al. 2011; Sillanpaa et al. 2012), animal (Zhao et al. 2004a, Xiong et al. 2011) and human (Li et al. 2009), and its application can be extended to genetic dissection of developmental process, including growth trajectory and allometric scaling (Ma et al. 2003, Li 2014), phenotypic plasticity based on gene-environment interaction (Wang et al. 2013b), drug response (Wang et al 2013b), and morphological shape (Fu et al. 2013). Recently, the integration of functional mapping and differential equations (Fu et al. 2011; Wang et al 2013b) have also been applied to widely emergent applications of dynamic systems.

The *Funmap2* package is developed to identify QTLs for a longitudinal trait based on functional mapping. It is implemented as a package for the freely accessible statistical software R. *Funmap2* implements a complete pipeline which includes data loading, QTLs scanning, significance value computing and significant QTLs reporting. The essence of functional mapping lies on the longitudinal curve of genetic effects and the covariance matrix that characterize longitudinal relationship of the trait. Although a logistic curve may describe the genetic effects in most of biological processes, that of reaction norm in continuously varying environment problem may not follow sigmoid shape. To address this, *Funmap2* provides sigmoid, Legendre, Pharmacology as the built-in resource, and it also allows users to customize the curve equation. Furthermore, to increase the statistical power, the covariance matrix can be chosen from several covariance structures used in IBM SPSS software, such as autoregressive, ante-dependence, or autoregressive moving average (Li et al 2010a). Considering the difficulty to know which combination of curve shape and covariance structure is the best for a longitudinal trait beforehand, we enable *Funmap2* to automatically choose the best curve shape and covariance structure from the built-in resource based on information criterion.

In addition to statistical analysis, *Funmap2* generates a publication-level report that visualize all results, including phenotype traits, QTL profile, significant QTL curve, and the permutation result. Additionally, the package provides a simulation module for testing the performance and demonstrating the use of the *Funmap2* on data generated by different models. Empirically, we observe the robust performance of *Funmap2* on simulated data with inheritability greater than 5%.

To the best of our knowledge, *Funmap2* is the only QTL mapping tool available for the longitudinal traits with open source software license. It is publicly available under an open-source software license: https://github.com/wzhy2000/Funmap2. In the following sections, we give a brief review of functional mapping in term of its statistical model. Then we focus on the detailed workflow of *Funmap2*. Lastly, we provide one example with codes and figures.

## Statistical Methods

Functional mapping is a statistical framework aiming at identifying QTLs that are significantly associated with a longitudinal phenotype of interest on an experiment population, such as recombinant inbred lines (RIL) and doubled-haploid (DH) population. Functional mapping computes maximum likelihood estimation (MLE) of mixture models that integrate the likelihood over QTL genotypes. The model also accounts for the 1) longitudinal trend of the trait using a continuous curve, i.e. growth trajectory for time-dependent traits and reaction norm for environment-dependent traits, and 2) internal correlation of traits across longitudinal measurement using a covariance matrix.

The model assumes that the longitudinal trend of the trait follows a particular mathematic curve. In statistical setting, the phenotypes of the trait at all time points follow a multivariate normal distribution, which therefore can be described by the following equation (1):

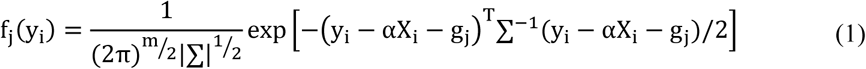

Assuming that y_i_ is a vector of measured values for individual *i* at *m* time points, which describes the phenotypic values; g_j_ denotes the overall mean vector for genotype *j*, and is described as a mathematic curve, X_i_ denotes the covariate for individual *i*, α is a vector of coefficient value for each covariate, ∑ is the covariance matrix. Therefore, f_j_(y_i_) is probability density function relating the measured traits of individual *i* to the combination of the genetic effects contributed by genotype *j* and covariates effects of individual *i.*

The estimated likelihood of genotype *j* for individual *i* are summed, weighted by corresponding conditional probability of QTL genotype given the adjacent marker and inbred type. The functional mapping method hence formulates the likelihood calculation of the mixture model as:

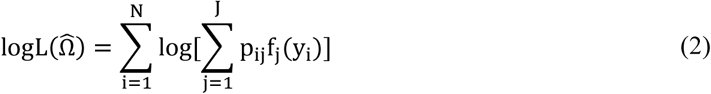

where each element p_ij_ indicating the genotypic possibility of subject *i* for gene *j* (QQ, Qq or qq), N denoting the individual number in the experiment population, Ω̂ denoting the random variables in this log likelihood function including covariate coefficient, curve parameters for multiple genotype, and covariance parameters. In equation (2), we prefer to use log likelihood function.

The likelihood function for the null hypothesis model, in which QTL does not affect the trait, is built as follows,

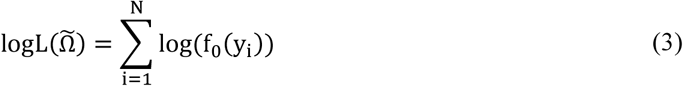

where f_O_(y_i_) differs from (1) in g_j_ by assuming the same longitudinal curve for all genotypes. Ω̃ denotes the estimated parameters in this log likelihood function.

The goal of functional mapping is to compute log likelihood ratio (LR) for each QTL, defined by the following equation, and then to choose the significant QTL with high value of LR.

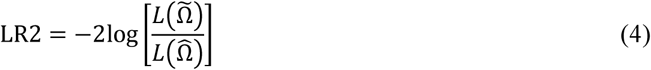

Intuitively, the null hypothesis H_0_ is that there is no gene that controls the growth process, and the alternative hypothesis H_1_ is the growth processes are different across the QTL genotypes. Where Ω̃ and Ω̂ are the maximum likelihood estimates of parameters under the hypothesis H_0_ and H_1_ respectively.

To do log-likelihood ratio test on complicated statistical models like the one used in functional mapping, permutation test is usually used to derive and compare against the null distribution. Whereas permutation test is generally applicable to various models, it is computationally intensive to implement. To address this, we propose a new approach called filtering method, to improve the computational efficiency of a permutation test. We first quantify the correlation between QTL and longitudinal data, using a genotype-oriented curve clustering method. Then, the QTLs which are highly correlated with the outcome were computed in the improved permutation tests (Wang et al, 2017). As a result, it significantly reduces the amount of computation in permutation tests and speeds up the computation for data analysis in functional mapping.

## Package Workflow

The *Funmap2* package is an open source package for R with automated data analysis for identifying the significant QTLs for longitudinal traits measured. The *Funmap2* pipeline includes modules for data import, curve fitting, QTL scanning, MLE computation, hypothesis testing, and data visualization. Figure 1 shows the workflow of *Funmap2*, with the left column being the analysis steps and the right column being the corresponding output obtained in each step.

**Figure 1:**
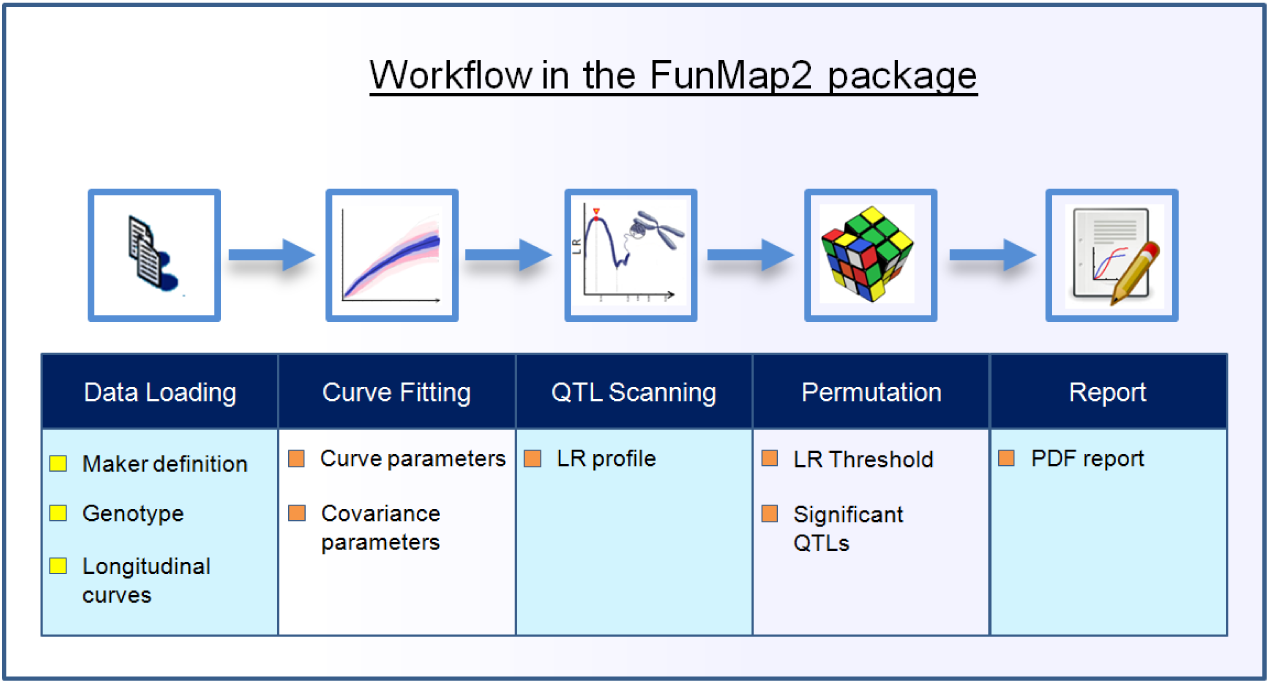
Workflow

User may either run the entire *Funmap2* workflow in one function call or step-by-step with customization. Often it is more convenient to call the main function *FM2.pipe* which automatically implements all tasks and outputs the summary information and figures in PDF format. In this function, sub-modules are called successively, functions involved include data loading (*FM2.load.data*), data estimation (*FM2.estimate.data*) for curve fitting and covariance selection, QTL scanning (*FM2.qtlscan*), permutation (*FM.permutation)*, and report generating (*FM2.report*). The typical calling is shown below.

~~~
# Call the pipeline in parallel computing.
r <- FM2.pipe(file.pheno.csv, NULL, file.geno.csv, file.marker.csv, “BC”, curve.type="logistic",
covar.type=“auto”, options=list(n.cores=10));
~~~

*Funmap2* requires users to provide experiment data and specifying several parameters. Phenotype file containing longitudinal traits, one genotype maker file for the experiment population, and one genetic marker information file are required. Covariate file is optional and is not provided in the above example. In addition to these experiment data, users need to specify the cross type of QTL mapping. 4 available types are provided in *Funmap2*, Backcrossing, F2, RIL and DH. Noted that in this automated run, user may either let *Funmap2* choose the optimal curve type and covariance structure or specify their values as functional arguments.

Alternatively, the data analysis can be conducted by the customizing workflow by successively running sub-modules of *Funmap2*, as illustrated below.

### 3.1 Data Loading

Input files for *Funmap2*, including the marker definition file, the genetic marker file, the phenotype file, and the covariate file, should be in CSV format. The function *FM2.load.data* reads these data files, checks the correctness of data format and the consistency of Individual IDs across all files and finally returns a data object wrapper in S3 class. Phenotypic traits, as well as curves describing longitudinal trend (if `curve.type` is specified) can be visualized by directly calling plot function on the returned S3 object.

### 3.2 Curve Fitting & Covariance Matrix Selection

The function *FM2.estimate.data* may be invoked to facilitate the manual selection of curve type and covariance structure. The most commonly used curve function is sigmoid, and is generally used to characterize growth curves of biological organisms. Curves other than sigmoid, such as those describing environment-dependent traits for which reaction norms are measured, are also provided. In total, *Funmap2* software implements 9 curves in the current version, including logistic, composite, and those generated by nonparametric method. When running the automated mode, i.e. curve type is not specified during the data loading, least square curve fitting followed by AIC (Akaike Information criterion) and BIC (Bayesian Information criterion) is used to determine the best curve type.

The internal correlation of traits measured at different longitudinal points are described using the covariance matrix, which is essential in the likelihood calculation. We included a comprehensive set of 13 covariance matrices including those employed by SPSS software and first order ante-dependence (Yap et al 2005). *Funmap2* also implements an automated way of selecting covariance matrix by AIC and MLE method. Users should be cautious of the cost of computational time when over-parametrizing the covariance matrix.

### 3.3 QTL Scanning

The function *FM2.qtlscan* estimates QTL effects as well as parameters that characterize longitudinal trend curve by scanning across all QTL positions. Specifically, the function conducts H_0_ and H_1_ hypothesis by MLE method and calculates LR2 values in Equation 4 at each QTL marker at every 1 cM position. The MLE method also outputs covariate coefficients, curve and parameter that optimizes the log likelihood. When finished, *FM2.qtlscan* by default generates the LR2 profile figure and highlight QTLs with the highest LR2 value on each chromosome. To determine the significant QTLs, user will need to run the permutation test (see the following section).

### 3.4 Permutation test

The distribution of log-likelihood ratio is difficult to be derived in analytical forms, especially for complicated distribution functions. To overcome this difficulty, a permutation test is generally used to obtain the null distribution and declare statistical significance of a QTL. One commonly encountered issue with permutation test, however, is the high cost of computational time, which becomes especially prominent when running on whole genome scale. The function *FM.permutation* includes two methods to address this issue. The first option is parallelization the computation in unix-based operating systems provided that multiple processor threads are available. The second one is to apply filtering method to reduce the computational intensity. This is done by pre-selecting the candidate QTLs with high QTL-trait correlation.

### 3.5 Report

The resulting objects returned by individual functions contain summarizing information (by the summary function), and can be visualized by the plot function. In addition, *Fumap2* includes the function *FM2.report* that can automatically generate reports, including tiled/overlapping curves describing the longitudinal trait, LR profile for all chromosomes, LR2 profile for QTL position and the curve for QTL position.

## Results

We use a data set from poplar as an example. The data are composed of 90 backcross individuals with 22 linkage groups and 275 molecular markers. The phenotypic values were measures throughout 11 years. The code for analyzing the data is shown below, and the results for QTL mapping are plotted in Figure 2 and Figure 3.

**Figure 2:**
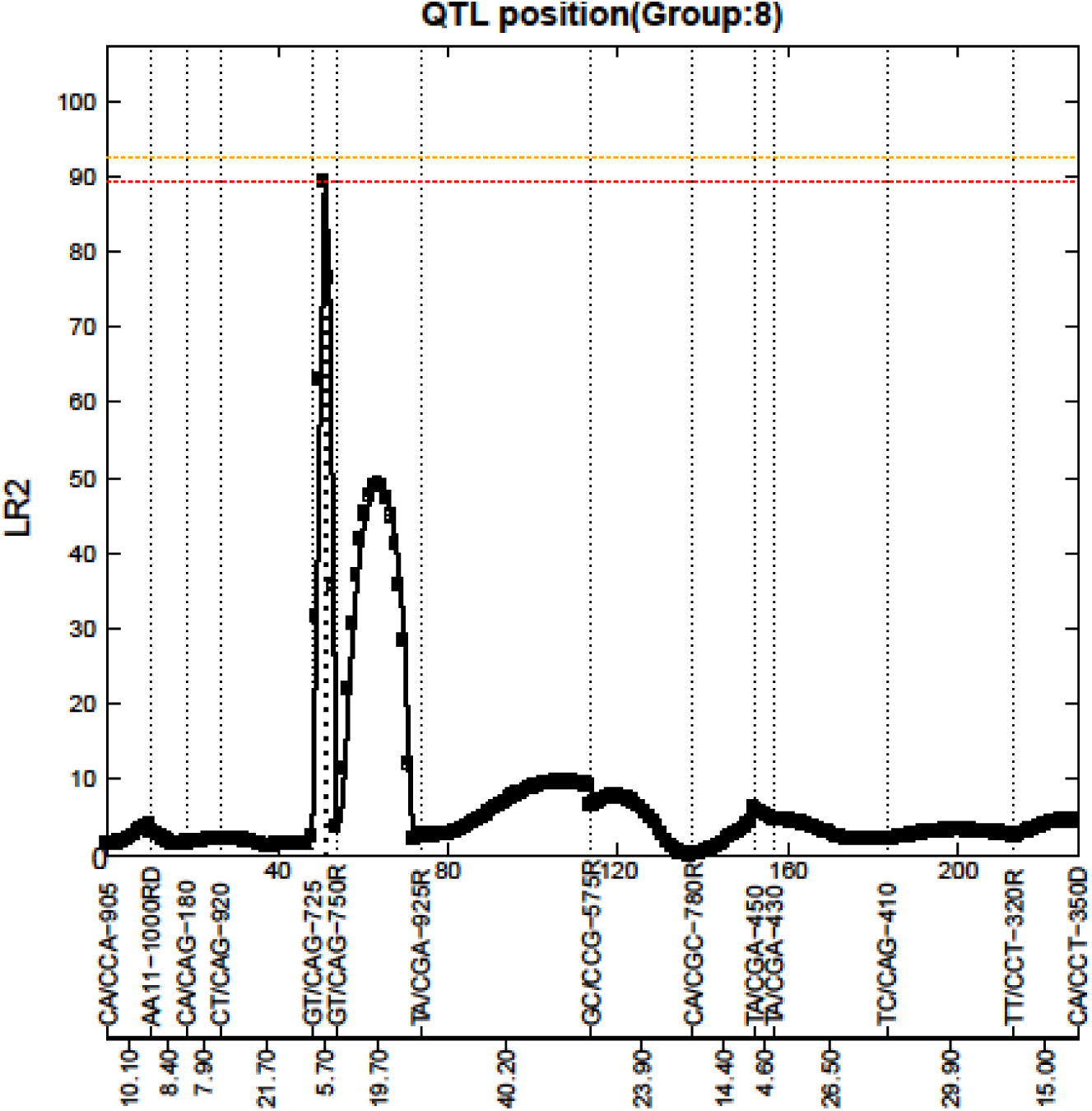
LR profile for QTL position

Figure 2. Profile of the likelihood ratio (LR) for one chromosome with 14 markers, with their names and genetic distance for each interval highlighted at the bottom. The peak with LR2=89.47 at interval [GT/CAG-725—GT/CAG-750R] is discernible at 3 cM, suggesting a potential loci for significant QTL.

**Figure 3:**
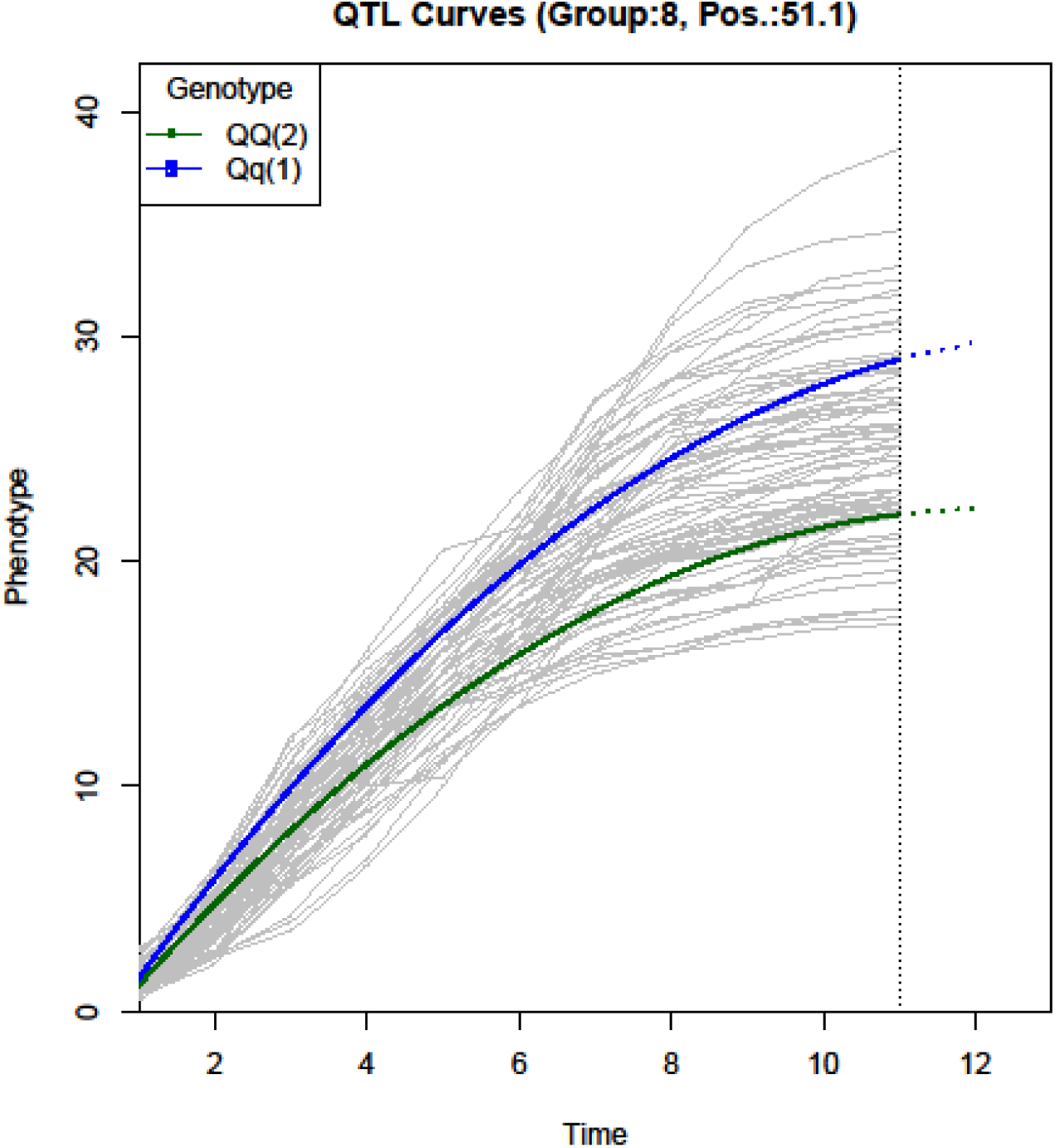
Growth curves for QTL position

Figure 3. Growth curve for QTL selected from figure 2. Logistic curve is chosen by curve fitting. The curve is colored by genotype. Noted that two genotypes show similar growth at early stages, but diverge at later stages, suggesting a QTL associated with growth kinetics.

~~~
# Load the pre-installed data for the example
file.pheno.csv <- system.file(“extdata”,“populus.BC.pheno.csv”, package=“Funmap2”)
file.geno.csv <- system.file(“extdata”,“populus.BC.geno.csv”, package=“Funmap2”)
file.marker.csv <- system.file(“extdata”,“populus.BC.marker.csv”, package=“Funmap2”)
r <- FM2.pipe(file.pheno.csv, NULL, file.geno.csv, file.marker.csv, “BC”, curve.type=“auto”, covar.type=“AR1”, options=list (n.cores=10));
~~~

## Discussion and Conclusion

Functional mapping model assumes the longitudinal traits follow a parametric or non-parametric curve, such as growth trajectory, Legendre polynomial, B-Spline (Yang et al 2009). Under this assumption, likelihood ratio and QTL effects derived from the parameters of parametric or non-parametric curve, are calculated by MLE function over all linkage groups. *Funmap2* implements the functional mapping framework with 9 curves and 13 covariance structures. Importantly, any new curve functions not implemented by *Funmap2*, can be easily imported into the package and assembled into the framework of MLE. It is open architecture for any biological curve to fit the longitudinal traits.

The longitudinal traits tend to strongly correlate between time points (time-dependent) or reaction norms (environment-dependent). Function mapping models this internal relation using covariance matrix, which may increase the statistical power for QTL detection. Whereas previous publications on functional mapping recommended the use of the most parsimonious covariance matrix (Yap et al, 2009), such as autoregressive, ante-dependence or autoregressive moving average (Li et al, 2010a), *Funmap2* also provides other covariance matrices implemented in IBM SPSS software, such as Compound Symmetry, Factor Analytic, Huynh-Feldt, Toeplitz. This is because although parsimonious covariance matrix can be computationally efficient, non-parsimonious covariance structures contain more parameters, and hence richer structures, which may potentially lead to better data fitting while minimizing the pitfall of overfitting when guided by information criterion.

Studies of QTL mapping for longitudinal traits other than functional mapping are unexpectedly rare, compared to that for QTL mapping on trait measured at a single point. The works include Yang et al 2006 and Funqtl (Kwak et al 2016). As a result, researches on the genetic basis underlying biological development and gene/ environment interaction are greatly limited. *Funmap2* provides a user-friendly way to dissecting these problems, and facilitates the building of precise genotype-phenotype relation model through QTL mapping. Besides mapped QTLs, estimates from curve functions may also provide insights for the understanding of the genetic, biochemical and physiological pathways governing developmental change (Wang et al. 2012). We are making the endeavor to develop a GUI version *Funmap2* to further facilitate the user to extract and interpret the data. At present, *Funmap2* supports experimental populations derived from a cross between two inbred lines, and is limited to four types, F2, backcross, recombinant inbred lines and double-haploid populations. Future versions of *Funmap2* shall be able to accommodate populations of more diverse structures and even multiple traits, epistasis effects, allometric, and QTL-QTL interaction, as functional mapping itself is rapidly evolving and is quickly merging with these fields.

## Funding

This study was supported by National Training Program of Innovation and Entrepreneurship for Undergraduates (201510022069), the Fundamental Research Funds for the Central Universities (BLX2013026), NSFC Grant (31470675).

